# CNLLRR: A Novel Low-Rank Representation Method for Single-cell RNA-seq Data Analysis

**DOI:** 10.1101/818062

**Authors:** Na Yu, Jin-Xing Liu, Ying-Lian Gao, Chun-Hou Zheng, Junliang Shang, Hongmin Cai

**Affiliations:** School of Information Science and Engineering, Qufu Normal University, Rizhao, Shandong, China; Qufu Normal University Library, Qufu Normal University, Rizhao, Shandong, China; Co-Innovation Center for Information Supply and Assurance Technology, Anhui University, Hefei, Anhui, China; School of Computer Science and Engineering, South China University of Technology, Guangzhou, Guangdong, China

## Abstract

The development of single-cell RNA-sequencing (scRNA-seq) technology has enabled the measurement of gene expression in individual cells. This provides an unprecedented opportunity to explore the biological mechanisms at the cellular level. However, existing scRNA-seq analysis methods are susceptible to noise and outliers or ignore the manifold structure inherent in the data. In this paper, a novel method called Cauchy non-negative Laplacian regularized low-rank representation (CNLLRR) is proposed to alleviate the above problem. Specifically, we employ the Cauchy loss function (CLF) instead of the conventional norm constraints in the noise matrix of CNLLRR, which will enhance the robustness of the method. In addition, graph regularization term is applied to the objective function, which can capture the paired geometric relationships between cells. Then, alternating direction method of multipliers (ADMM) is adopted to solve the optimization problem of CNLLRR. Finally, extensive experiments on scRNA-seq data reveal that the proposed CNLLRR method outperforms other state-of-the-art methods for cell clustering, cell visualization and prioritization of gene markers. CNLLRR contributes to understand the heterogeneity between cell populations in complex biological systems.

**Author summary:** Analysis of single-cell data can help to further study the heterogeneity and complexity of cell populations. The current analysis methods are mainly to learn the similarity between cells and cells. Then they use the clustering algorithm to perform cell clustering or downstream analysis on the obtained similarity matrix. Therefore, constructing accurate cell-to-cell similarity is crucial for single-cell data analysis. In this paper, we design a novel Cauchy non-negative Laplacian regularized low-rank representation (CNLLRR) method to get a better similarity matrix. Specifically, Cauchy loss function (CLF) constraint is applied to punish noise matrix, which will improve the robustness of CNLLRR to noise and outliers. Moreover, graph regularization term is applied to the objective function, which will effectively encode the local manifold information of the data. Further, these will guarantee the quality of the cell-to-cell similarity matrix learned. Finally, single-cell data analysis experiments show that our method is superior to other representative methods.

## Introduction

In recent years, a large number of single-cell RNA sequencing (scRNA-seq) data have been generated due to the development of next-generation sequencing technologies [1]. At the same time, scRNA-seq data contain a wealth of information on biological function and gene regulation. Analysis and research on these information pave us a way to observe individual cells unprecedentedly [2]. It thus provides us possibilities to explore the heterogeneity and complexity of cell population. It also offers us an unprecedented opportunity to learn the biological mechanisms and functional diversity at the cellular level.

With the help of scRNA sequencing, identification of subpopulation of cells [3] is now possible. The identification can be considered as a clustering problem through the similarities of gene expression. Those cells with high similarity will be categorized into the same subgroup. Cell clustering contributes to the targeted therapy of disease and the discovery of new cell subtypes. By using visualization methods, such as *t*-distributed stochastic neighbor embedding (t-SNE) [5], biologists can intuitively understand the distribution of the cells. Through prioritizing the genes that define the subgroup of cells, a new direction is provided for us to study cell differences.

Currently, many methods for single-cell clustering have been proposed [6, 7]. Generally speaking, these methods are mainly to learn the similarity between cells and cells. Then they use the clustering algorithm to perform cell clustering or downstream analysis on the obtained similarity matrix. Therefore, constructing accurate cell-to-cell similarity is crucial for scRNA-seq data analysis. Up to now, many efforts have been taken, for example, Xu et al. used the idea of shared nearest neighbors to measure the similarity between cells [8]. To enhance clustering accuracy, Wang et al. proposed the single-cell interpretation via multi-kernel learning (SIMLR) [9], which obtains a similarity matrix by learning the appropriate distance metric. Then Kim et al. designed a new framework based on improved SIMLR by using Pearson correlation as the measure of similarity [10]. Influenced by the assumption that neighbor information can measure the similarity, Jiang et al. proposed a method called Corr, which is based on differentiability correlation of cell pairs [11].

The aforementioned method relied on the accurate pairwise similarities among the cells. However, the similarity prones to be corrupted by noises or outliers. The high dimensional characteristics of the scRNA data further deteriorates the accuracy of the similarity calculation by involving an averaging operation. Alternatively, recent advances focus on capturing the global structure of the data. Zheng et al. proposed a subspace clustering algorithm called SinNLRR based on low-rank representation (LRR) to identify cell types [4]. Although the SinNLRR method achieves satisfactory experimental results, two issues still need to be considered. One is that single-cell data inevitably contain noise (for example, in the process of measuring and collecting scRNA-seq data) and outliers. However, the noise matrix of the SinNLRR method is constrained by the Frobenius norm, in which case its performance is affected. In addition, the common *L*_1_-norm, *L*_2,1_-norm or Frobenius norm is applied to the noise term, which can only model specific noise. In general, *L*_1_-norm can model the noise that matches the Laplacian distribution, *L*_2,1_-norm constraint is imposed to tackle sample-specific outliers, and Frobenius norm can well characterize Gaussian noise [12]. Unfortunately, noise and outliers are generally unknown and have complex statistical distributions. Therefore, a reasonable new strategy is needed to effectively deal with noise and outliers. Another issue is that there are low-dimensional manifold structures in the high-dimensional sample space of data. If the manifold information is not considered, the similarity matrix learned will not better reflect the similarity among cells. This will seriously affect the results of single-cell analysis.

To overcome the above limitations, we design a novel Cauchy non-negative Laplacian regularized low-rank representation (CNLLRR) method for clustering cells, visualizing cells and prioritizing gene markers. Specifically, Cauchy loss function (CLF) constraint is applied to punish noise matrix. It can effectively reduce the influence of a single sample containing noise and outliers and avoid to be affected pronouncedly when the sample contains large noise. In addition, the influence function of CLF has an upper limit when estimating residuals, so it seldom relies on the distribution of noise and outliers [13]. Therefore, the CNLLRR method is robust to noise and outliers. Moreover, to preserve the intrinsic geometric local information in scRNA-seq data, the graph regularization term is applied to the objective function. The main contributions of this paper are as follows:

1. A novel method called CNLLRR is presented to capture the global structure of the data. CNLLRR uses the CLF instead of the conventional norm constraints on the noise term, which effectively reduces the influence of a single sample. Therefore, the robustness of the CNLLRR method to noise and outliers is improved. Moreover, the CNLLRR method also applies graph regularization terms to encode manifold information of the data. These will improve the quality of the learned similarity matrix.
2. The alternating direction method of multipliers (ADMM) is employed to optimize the CNLLRR method. And the robustness of our method is proved in theory and experiments.
3. We perform comprehensive experiments to validate the CNLLRR method on the scRNA-seq data. The experimental results show that our method is meaningful and superior to several representative methods.

The rest of this paper is organized as follows. In Materials and methods Section, LRR, CLF, and graph regularization are simply reviewed. The CNLLRR method and its optimization are also described in detail. In Results and discussion Section, the experimental results on scRNA-seq data are demonstrated. The paper is concluded in Conclusion Section.

## Materials and methods

### Low-rank representation

LRR is a widely used data analysis technique for exploring multiple low-dimensional subspace structures [14, 15]. In the field of bioinformatics, a scRNA-seq dataset can be represented by a matrix. The columns and rows of the matrix represent cells and the level of gene expression in different cells, respectively.

Given a data matrix **X** = [**x**_1_, **x**_2_, *…*, **x**_*n*_] ∈ *R*^*m*×*n*^, where **x**_*j*_ can be considered as a linear combination of the bases in the dictionary [16]. Therefore, the data matrix plays the same role as the dictionary which helps to discover the potential affinity between data samples. LRR decomposes the data matrix into the low-rank representation of the data and the noise, the purpose of which is to search the lowest rank that can represent all the data [17]. Therefore, the objective function of the LRR can be expressed as the following rank minimization problem:

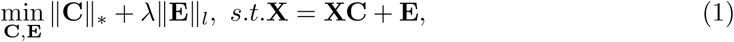

where **C** ∈ *R*^*n*×*n*^ and **E** ∈ *R*^*m*×*n*^ represent the coefficient and the noise matrices, respectively. ‖ · ‖_*_ is the nuclear norm of the matrix, which is defined as the sum of the singular values of the matrix. And ‖ ·‖_*l*_ is some sort of regularization strategy, such as ‖ ·‖_1_, ‖ ·‖_2,1_, and ‖ · ‖_*F*_. *λ* ≥ 0 denotes the weighting parameter which is used to balance the two terms.

Recently, Zheng et al. proposed a modified version of LRR to detect cell types in scRNA-seq data [4]. If the expressions of cells lie in the same subspace, it is implied that these cells are most likely of the same type. Substituting **X** − **XC** for **E**, the objective function of SinNLRR is defined as

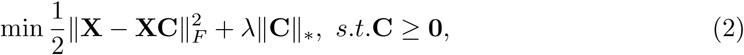

where ‖ · ‖_*F*_ represents the Frobenius norm of matrix. **C** can be seen as a representation of **X**, with **C** ≥ **0** meaning the non-negative similarity between cells. The penalty is adaptive due to the selection of *λ*.

### Cauchy loss function (CLF)

In real-world applications, noise and outliers are common in the data. What is more, they are usually unknown and complex. Therefore, a robust loss function is urgently needed to deal with this problem. Recently, CLF, as a robust estimator, has been successfully used in face recognition [18] and image clustering [13]. CLF can also be applied in solving the minimum issue in the form of the following optimization problem:

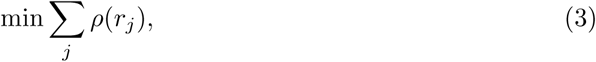

where *r*_*j*_ is the residual between the *j*-th data and its estimated value. *ρ*(·) represents a symmetric and positive-definite function where there is a unique minimum at zero. The *ρ*(·) function in CLF is log (1 + (*x/a*)^2^), hence the influence function of it can be calculated as

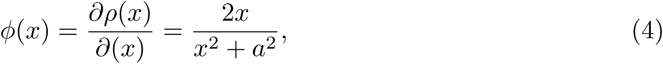

where *a* is a constant. *ϕ*(·) reflects the impact of sample points changes on the parameter estimation. It can be seen from Eq (4) that as the residual *x* increases, the influence function of CLF changes slightly. In other words, the influence function of the CLF has an upper bound when estimating the residual [13]. Therefore, using CLF as a constraint on the noise matrix in the model helps to improve the robustness of the method.

### Graph regularization

Due to the rapid development of manifold learning, people pay more and more attention to the inherent nonlinear structure of data. Graph regularization is based on local invariance assumption that if two sample points are close to each other in the high-dimensional data space, they are also close in the new low-dimensional space [19]. Hence graph regularization can capture the potential local manifold structure in the original data. For a given data matrix **X**, we use the data points as the vertices to construct a *k*-nearest neighbor (*k*-NN) graph [20]. Then, a symmetric weight matrix **W** is defined [21], whose element **W**_*ij*_ represents the weight of edge that joins the vertex **x**_*i*_ and the vertex **x**_*j*_. It can be calculated as

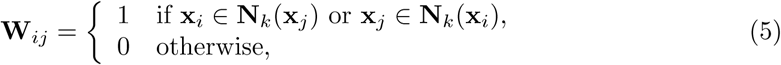

where **N**_*k*_(**x**_*i*_) represents the set of *k* nearest neighbors of **x**_*i*_. In mathematics, the graph regularization can be measured by

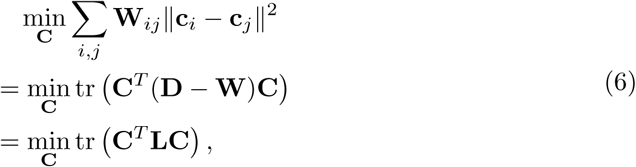

where **c**_*i*_ and **c**_*j*_ denote the mappings of **x**_*i*_ and **x**_*j*_ under some tranformations [22]. The degree matrix **D** is a diagonal matrix whose non-zero element **D**_*ii*_ = ∑_*j*_ **W**_*ij*_. **L** is the graph Laplacian matrix of **X** [23].

### Cauchy non-negative Laplacian regularized low-rank representation

In this section, we introduce the Cauchy non-negative Laplacian regularized low-rank representation (CNLLRR) method in detail. The CNLLRR algorithm is summarized in Algorithm 1.

#### Objective function

LRR has been successfully applied in many fields because it can effectively explore the global structure of data [22]. However, data are inevitably contaminated by noise and outliers in real world applications. To alleviate this issue, some methods are proposed as the variants of LRR by adding a regularization on the noise matrix. Nevertheless, noise and outliers are usually unknown and have complex statistical distributions. The *L*_1_-norm, the *L*_2,1_-norm and the Frobenius norm constraints are no longer suitable for most real world situations. Meanwhile, it is necessary to consider the low-dimensional manifold structure embedded in high-dimensional space in LRR-based methods.

To overcome the above limitations, a method called CNLLRR is proposed. CLF rather than conventional norm constraints is applied to penalize the noise matrix. CLF can effectively reduce the influence of a single sample containing noise and outliers. Moreover, it seldom relies on the distribution of noise and outliers [13] which helps to improve the robustness of the method. In addition, a graph regularization term is applied to encode the potential manifold information of data. In conclusion, the formulation of CNLLRR can be described as follows:

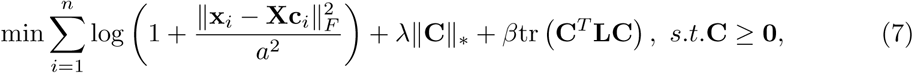

where *λ* and *β* denote penalty factors for balancing the regularization terms. **c**_*i*_ is a representation of the *i*-th data **x**_*i*_. Similar to the SinNLRR method, we replace **X** − **XC** with the noise matrix **E**. The CLF constraint is applied on it. To achieve the computational separability of Eq (7), an auxiliary matrix **J** ∈ *R*^*n*×*n*^ is introduced. Then CNLLRR can be rewritten as the following optimization problem:

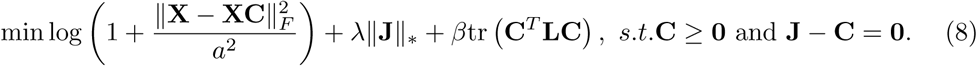

#### Optimization of CNLLRR

Alternating direction method of multipliers (ADMM) is used to optimize the CNLLRR method. Then the augmented Lagrangian function of Eq (8) can be defined as follows:

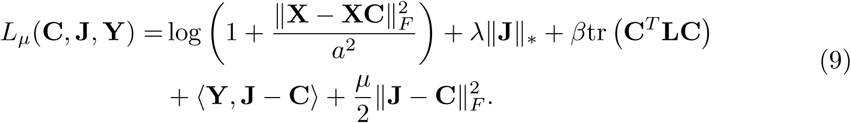

By using ⟨**A**, **B**⟩ = tr (**A**^*T*^ **B)**, Eq (9) can be further rewritten as

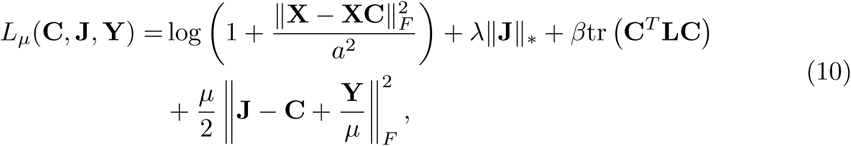

where **Y** ∈ *R*^*n*×*n*^ represents the Lagrange multiplier. *µ* denotes the user-defined parameter [4]. We can easily solve the problem Eq (10) by alternately updating one variable while fixing other variables.

First, **J**^*K*^ and **Y**^*K*^ are fixed when solving **C**^*K*+1^. And *L*_*µ*_(**C**, **J**^*K*^, **Y**^*K*^) is optimized, we get

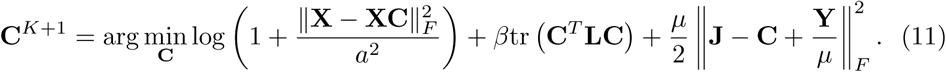

Through algebraic calculations, there exists an explicit solution of **C**,

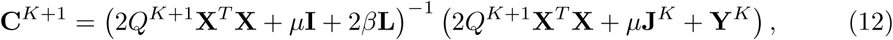

where 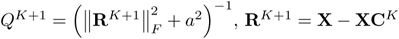 and **I** denotes an identity matrix.

Similarly, the subproblem used to update **J** can be calculated as

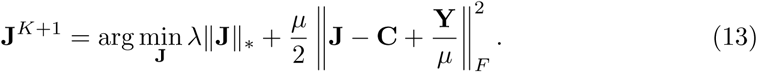

It is a quadratic optimization with low-rank constraint. It has an explicit solution by soft-thresholding,

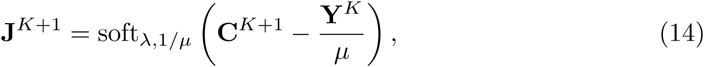

where soft_*λ*,1*/µ*_(·) represents the soft-thresholding operator [24]. And soft_*λ*,1*/µ*_(**A**) = **U***D*_*λ*,1*/µ*_ (**Σ**) **V**^*T*^, **A** = **UΣV**^*T*^. The element on the diagonal of matrix **Σ** is *σ*_*ii*_, *D*_*λ*,1*/µ*_ (**Σ**) = diag (max(*σ*_*ii*_ − *λ/µ*, 0)). Then the solution of **Y** is given by

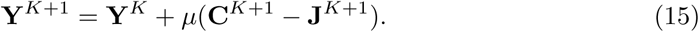

Finally, the optimization procedure for solving the proposed CNLLRR method is presented in Algorithm 1.

**Algorithm 1.**
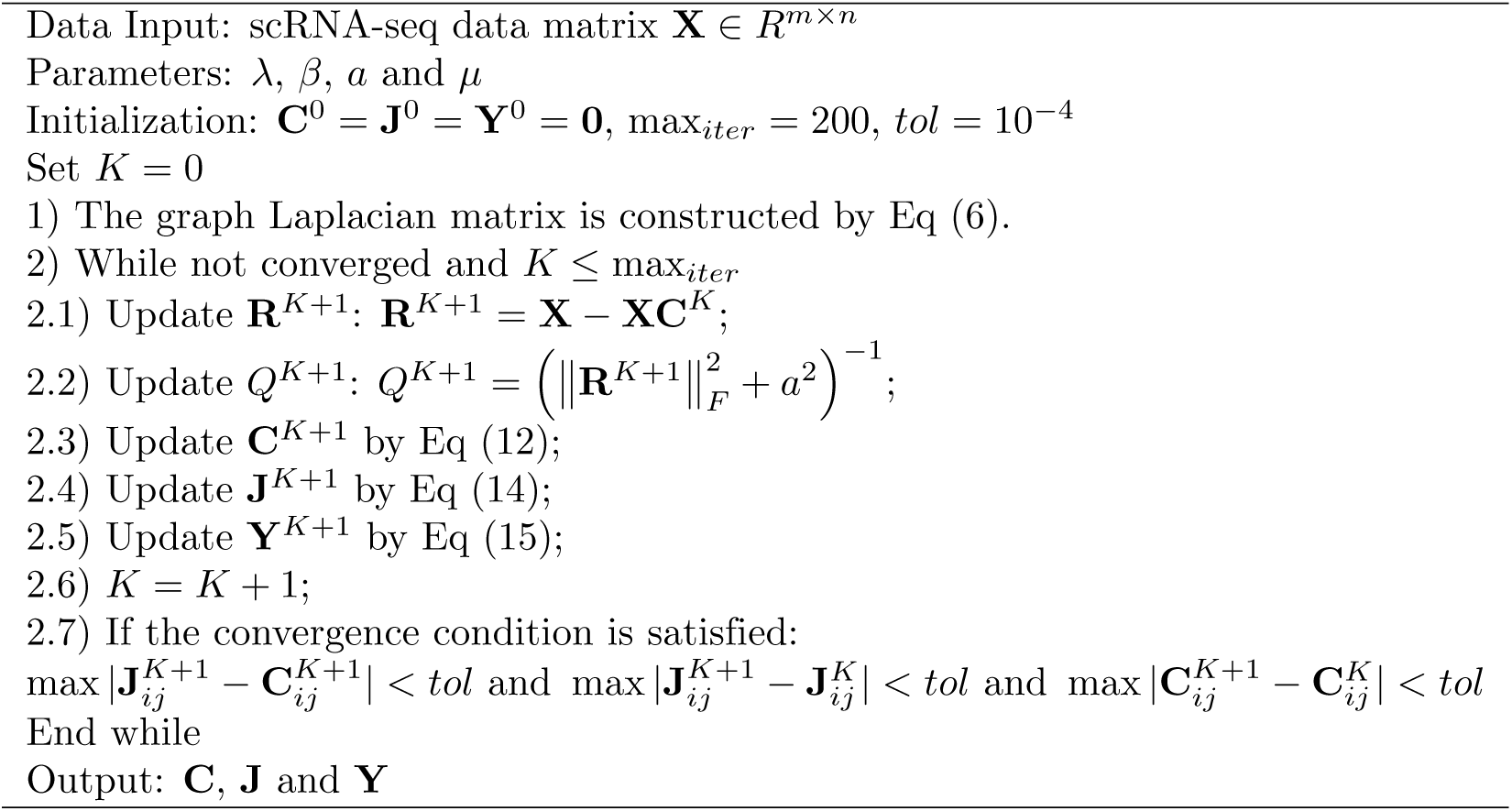
CNLLRR.

#### Robustness analysis

In this subsection, we analyze the robustness of the proposed CNLLRR method. From the viewpoint of statistics, using the robust estimator to measure errors is beneficial to enhance the robustness of the algorithm. Fig 1 shows the different estimators and corresponding influence functions. *L*_2_ estimator, *L*_1_ estimator and CLF are defined as *ρ*_2_(*x*) = *x*^2^, *ρ*_1_(*x*) = |*x*| and *ρ*_*c*_(*x*) = log (1 + (*x/a*)^2^), respectively. Their influence functions are represented as 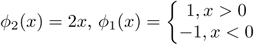 and *ϕ*_*c*_(*x*) = 2*x/x*^2^ + *a*^2^, respectively. The influence function can reflect the impact of sample points changes on parameter estimation. In Fig 1(b), the influence of a sample point on the parameter estimation increases linearly with respect to its error for *L*_2_ estimator. Therefore, *L*_2_ estimator is sensitive to noise and outliers. From Fig 1, we can also observe that *L*_1_ estimator can reduce the impact of large error on parameter estimation. However, with the error increasing, the influence function does not cut off. In Fig 1(b), for CLF, as the error increases, its influence function value tends to be zero. That is to say, the influence function of CLF has an upper limit. This is the reason why the CNLLRR method is robust to noise and outliers.

**Fig 1.**
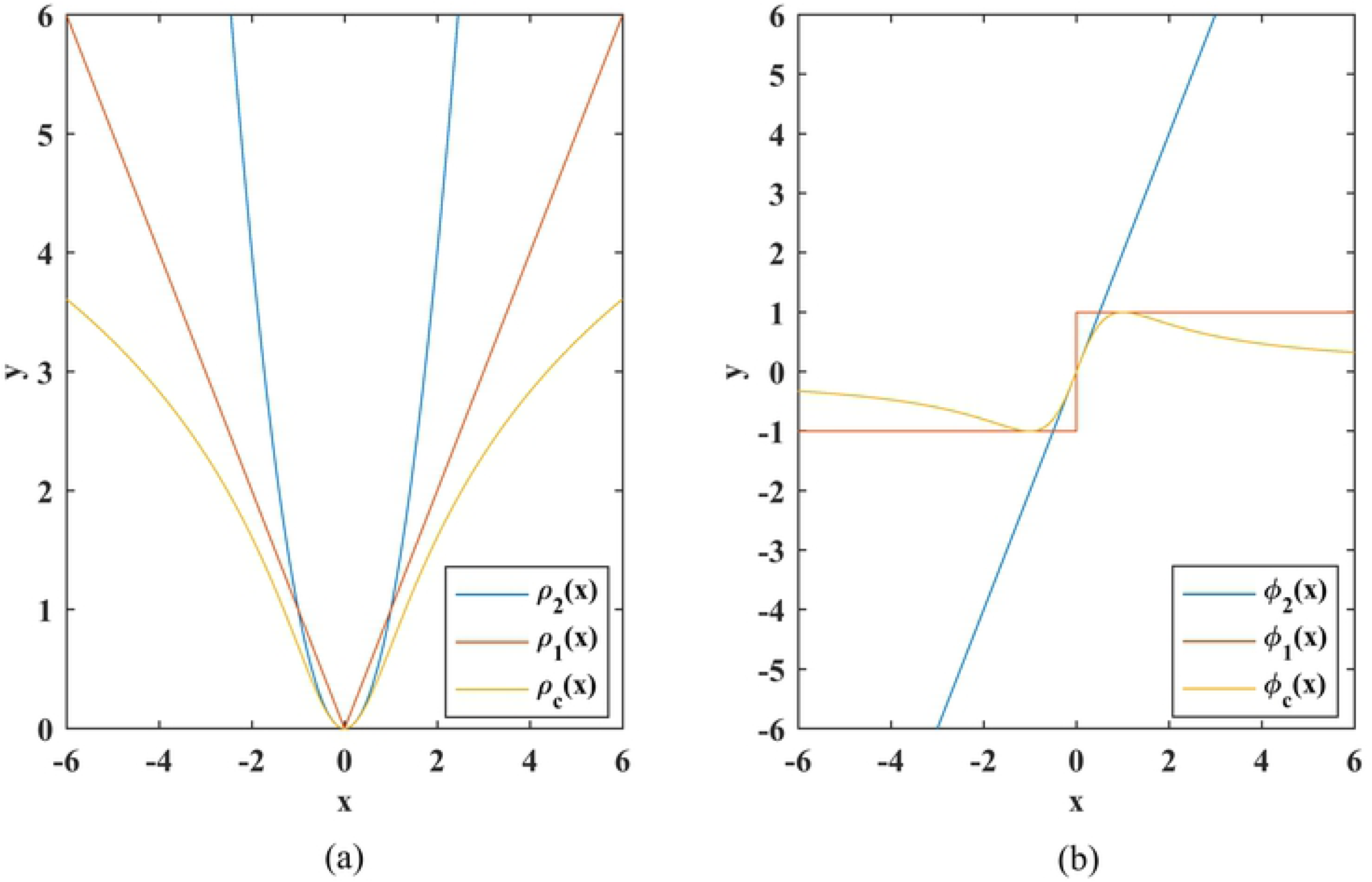
Explanation of the different estimators and corresponding influence functions. (a) Estimators. (b) Influence functions.

Next, we design an experiment to further illustrate the robustness of CNLLRR. Another method SinNLRR is also used for comparison. The experiment is conducted on a synthetic dataset consisting of 200 two-dimensional data points and the results are plotted in Fig 2. The subspace learning ability of the two methods on clean synthetic dataset can be revealed by Fig 2(a). From Fig 2(b) to 2(d), there are 20, 120, 150 randomly contaminated data points in the synthetic dataset, respectively. It can be seen that the subspace recovering ability of SinNLRR is severely limited with the number of contaminated points increasing. In contrast, CNLLRR can still explore the subspace structure successfully, even in the case of extreme noise points. In conclusion, our proposed CNLLRR method is more robust than SinNLRR.

**Fig 2.**
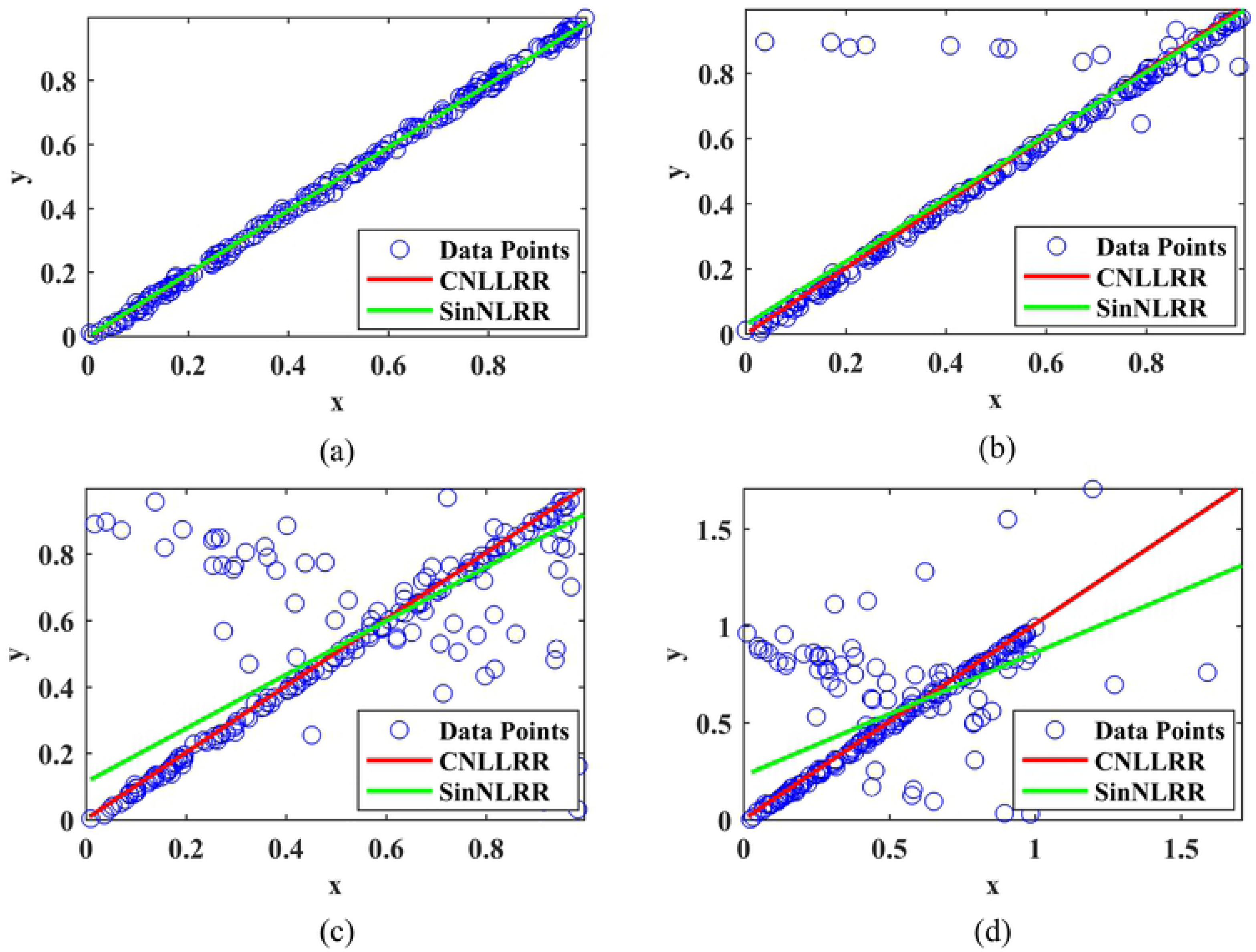
Robustness comparison of CNLLRR and SinNLRR on the synthetic dataset. (a) Clean dataset. (b) 20 contaminated data points. (c) 120 contaminated data points. (d) 150 contaminated data points.

#### Convergence and complexity analysis

We use the ADMM method to optimize the proposed CNLLRR method. As shown in Algorithm 1, **C**, **J**, and **Y** are iteratively updated until the convergence condition of CNLLRR is satisfied or the number of iterations exceeds the maximum we set. Therefore, the convergence of our method can be guaranteed. We also performed experiments to verify the convergence of the CNLLRR method. In Fig 3, the x-axis and y-axis represents the number of iterations and the error value, respectively. We can observe that as the number of iterations increases, the error value decreases gradually. This proves that the CNLLRR method has the property of fast convergence on all datasets. In addition, *O* notation is used to characterize the computational complexity. The computational cost of CNLLRR is mainly produced in the procedure of updating **C** and **J**. Futhermore, each time **C** is updated, **R** and *Q* also need to be calculated. Hence the time complexity of updating **C** can be calculated as *O* (*n*^3^ + *mn*^2^)according to Eq (12). As shown in Eq (14), singular value decomposition (SVD) needs to be employed when updating J whose computational cost is *O* (*n*^3^). Moreover, data graph also requires *O* (*mn*^2^) to be built. Assuming that *k* is the number of iterations of the algorithm, the total computational complexity of CNLLRR is *O* (*kn*^3^ + *kmn*^2^ + *mn*^2^).

**Fig 3.**
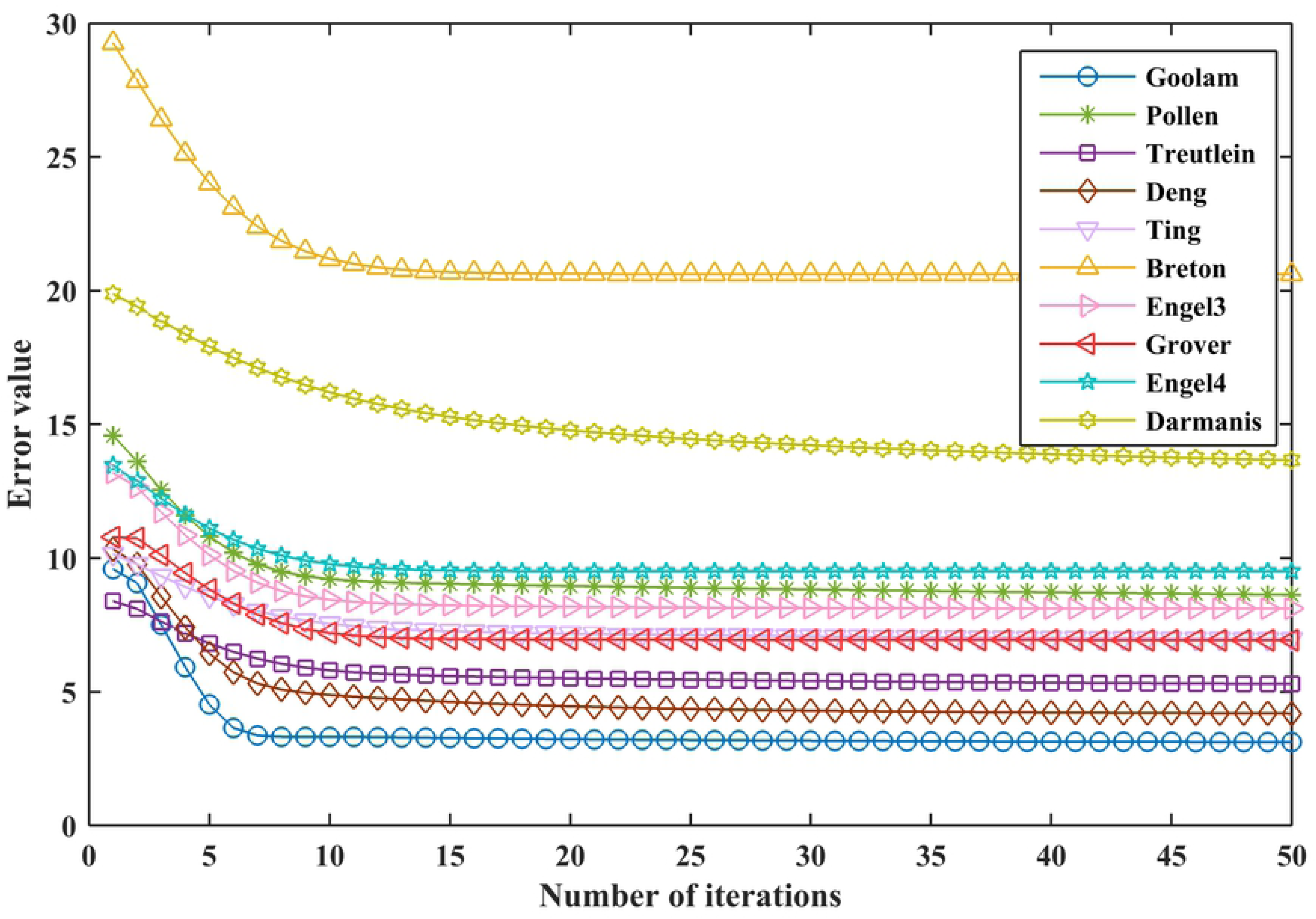
Convergence curves of CNLLRR on ten datasets.

#### Framework of CNLLRR

In this subsection, we use Fig 4 to describe the entire framework of CNLLRR in detail. Fig 4 can be seen as consisting of two main steps.

**Fig 4.**
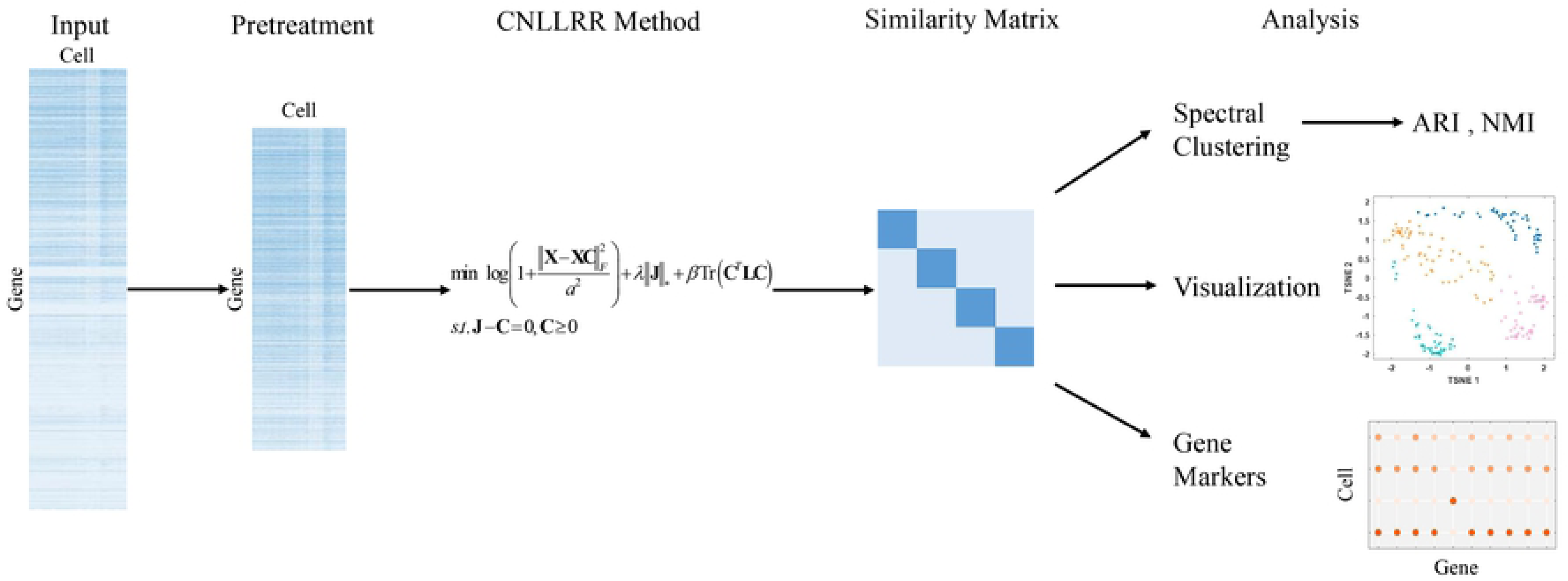
The entire framework of CNLLRR.

The first step is the adoption of the CNLLRR method. Specifically, preprocessing in needed for the input data matrix. The preprocessing includes gene filtering [25] and *L*_2_-norm normalization [26]. Gene filtering is commonly used in single-cell analysis,which aims to filter out the 5% genes that expresses less than all cells. *L*_2_-norm normalization normalizes each column vector of the matrix to 1, ie., 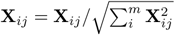. It can eliminate the scale differences between samples. After preprocessing, we use data matrix as the input of CNLLRR method to get the corresponding coefficient matrix **C**. Then we obtain the similarity matrix which has a better block diagonal structure in the same way as SinNLRR [4]. The localized similarity matrix is defined as

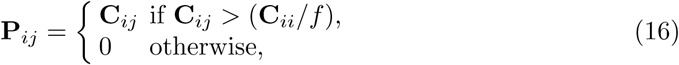

where *f* denotes the relaxation coefficient. Like SinNLRR, the value of *f* is also 1.5 in our method. The similarity matrix can be easily obtained through **S** = **P**^*T*^ + **P**.

The second step is the single-cell analysis. We perform spectral clustering on the learned similarity matrix. In addition, the similarity matrix is visualized and gene markers are prioritized.

## Results and discussion

In this section, clustering cells, visualizing cells and prioritizing gene markers are performed to analyze the performance of the CNLLRR method. In addition, t-SNE [5], K-means, PCA [27], sparse subspace clustering (SSC) [28], SIMLR [9], Corr [11], and SinNLRR [4] are used as comparison methods to verify the effectiveness of the proposed method.

### Datasets

With the launch of the Human Cell Atlas program, high-throughput single-cell sequencing technology has developed rapidly. This allows us to study gene expression in individual cells due to the large amount of valuable and unpredictable information from Single-cell data generated by sequencing technology. Studies in these information offer us a new insight into the learning of biological mechanisms at cellular level. In this paper, we analyze the performance of CNLLRR on scRNA-seq data, including Goolam [29], Pollen [30], Treutlein [31], Deng [32], Ting [33], Breton [34], Engel3 [35], Grover [36], Engel4 [35], and Darmanis [37]. The Engel dataset can be divided into 3 or 4 categories based on cell types, and they are represented as Engel3 and Engel4, respectively. In the experiment, gene filtering and *L*_2_-norm normalization are utilized to preprocess the data. Specific information on scRNA-seq data is listed in Table 1.

**Table 1.** Summary of ten scRNA-seq data.

| Datasets | Cells | Genes | Cell types | Species |
| --- | --- | --- | --- | --- |
| Goolam | 124 | 40315 | 5 | Mus musculus |
| Pollen | 249 | 14805 | 11 | Homo sapiens |
| Treutlein | 80 | 959 | 5 | Mus musculus |
| Deng | 135 | 12548 | 7 | Mus musculus |
| Ting | 114 | 14405 | 5 | Mus musculus |
| Breton | 957 | 20689 | 4 | Homo sapiens |
| Engel3 | 203 | 21690 | 3 | Homo sapiens |
| Grover | 135 | 14739 | 2 | Mus musculus |
| Engel4 | 203 | 23337 | 4 | Homo sapiens |
| Darmanis | 420 | 22085 | 8 | Homo sapiens |

### Parameter setting

In fact, the appropriate parameter setting will contribute to the performance of our method. There are four parameters in the proposed CNLLRR, which are the penalty factors *λ* and *β*, the constant *a*, the user-defined parameter *µ*. We use the grid search algorithm to select the optimal parameters. The values of *λ* and *β* are set in intervals [0.1, 1.3] and [0.001, 100], respectively. It should be noted that the value of *β* was changed in the Breton dataset from {3, 4, 5, 6, 7}. As for *a* and *µ*, the variation ranges of them are [0.001, 1] and [6, 11], respectively. Fig 5, Fig 6, and Fig 7 demonstrate the performance of CNLLRR with different *λ, β, a*, and *µ* on ten scRNA-seq datasets, respectively. From Fig 5 and Fig 6, we can observe that the proposed method is sensitive to parameters *λ* and *β*. Fortunately, *λ* and *β* can be selected to achieve better performance in the aforementioned interval. We can find in Fig 7(a) and 7(b) that as the values of *a* and *µ* increase, the behavior of CNLLRR is relatively stable. That is to say, the proposed method is robust to these two parameters. From Fig 7(b), we can observe that all ten datasets can achieve satisfactory clustering results. For convenience, we set the value of *µ* to 10 in the following experiment. In conclusion, reasonable parameter values can be selected from Fig 5 to Fig 7.

**Fig 5.**
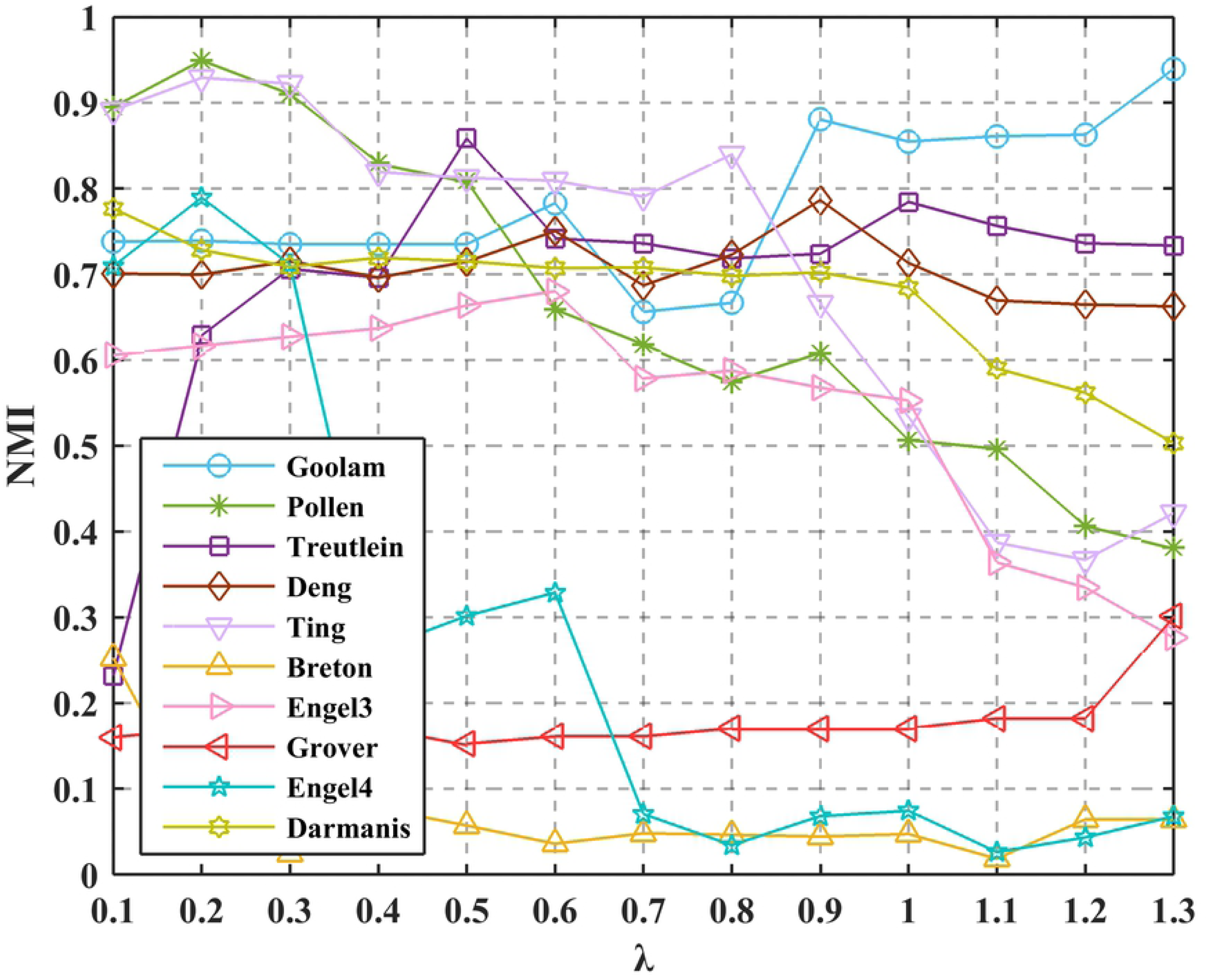
Performance of the CNLLRR set with different values of *λ*.

**Fig 6.**
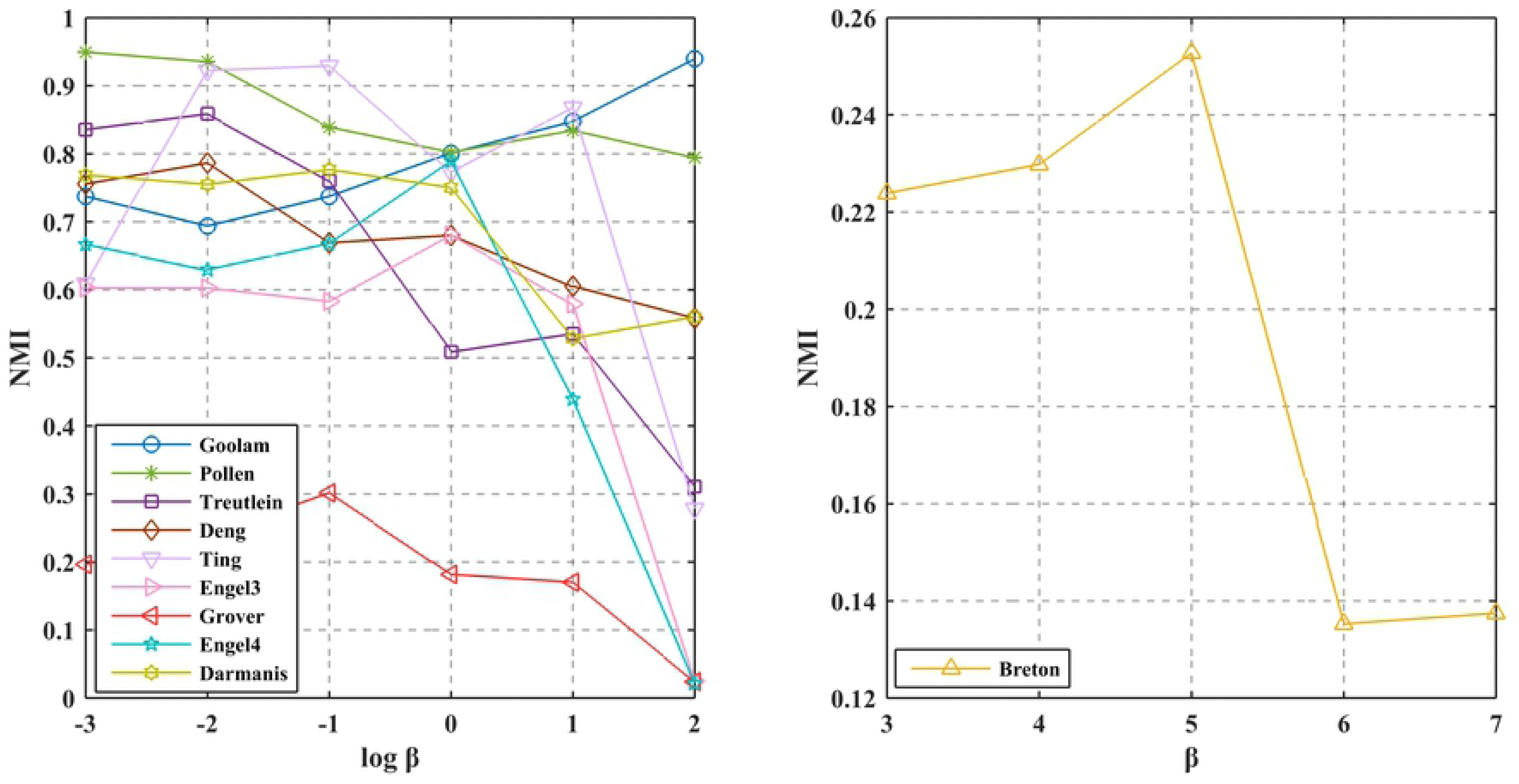
Performance of the CNLLRR set with different values of *β*.

**Fig 7.**
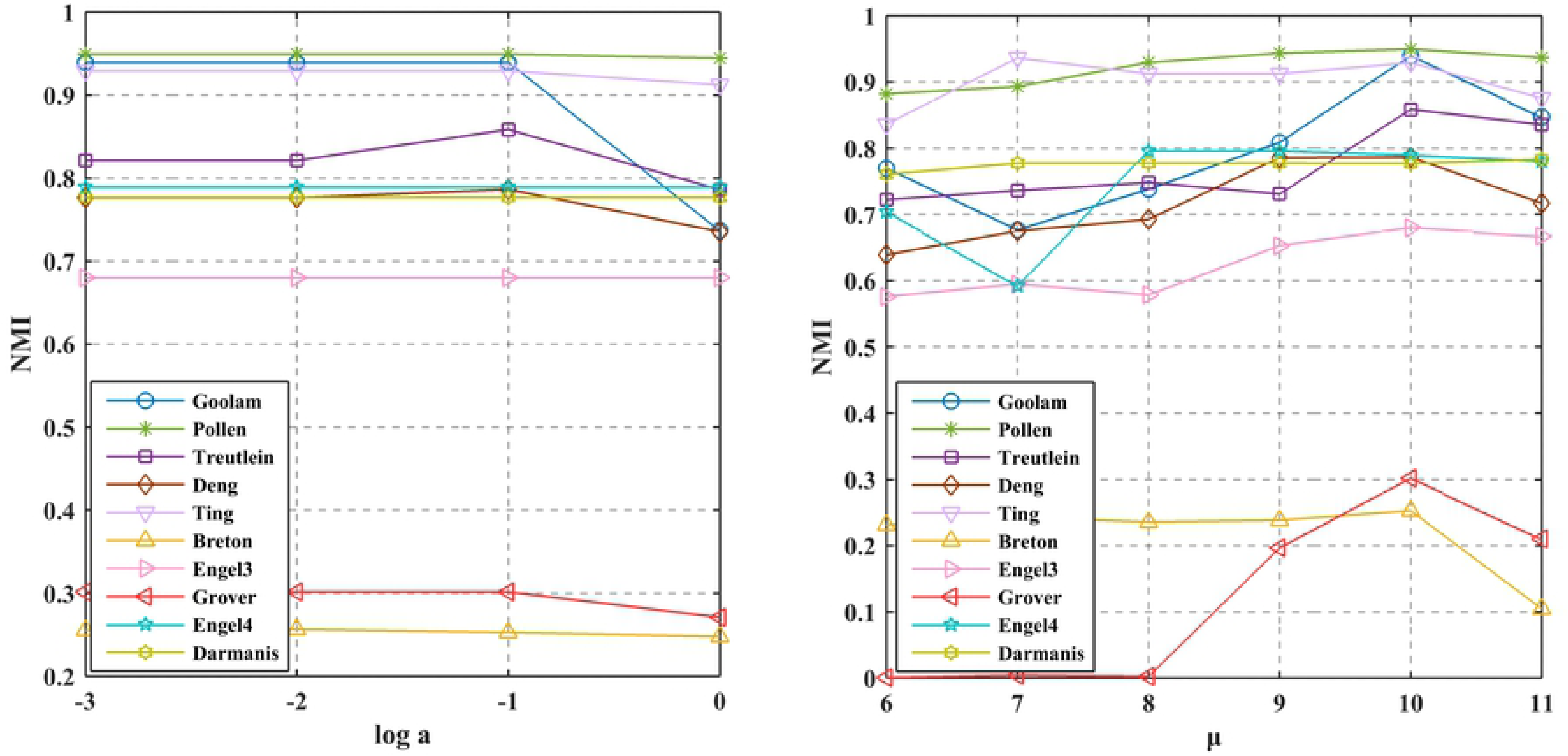
Performance of the CNLLRR versus parameters *a* and *µ*. (a) The different values of *a*. (b) The different values of *µ*.

### Clustering results

In this section, we perform clustering experiments to illustrate the effectiveness of the proposed method on scRNA-seq data. The spectral clustering algorithm is performed on the similarity matrix, which can divide cells into different subgroups according to cell types.

#### Evaluation metrics

To quantify the clustering results, we employ two widely used evaluation metrics: Adjusted Rand Index (ARI) [38] and Normalized Mutual Information (NMI) [39]. ARI is more suitable for sample-balanced datasets compared with NMI [40]. Let *T* = {*T*_1_, *T*_2_*…, T*_*K*_} and *P* = {*P*_1_, *P*_2_*…, P*_*K*_} represent the ground truth cluster set and the cluster set predicted by the clustering method, respectively. The ARI can be computed by

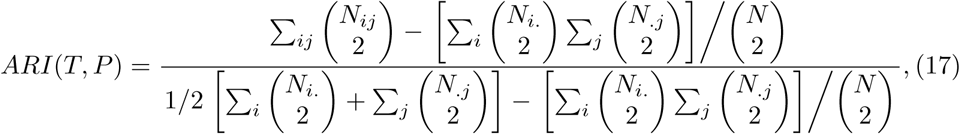

where *N*_*ij*_ represents the number of sample points in *T*_*i*_ and *P*_*j*_. The number of sample points in *T*_*i*_ is represented by *N*_*i.*_. And the number of sample points in *P*_*j*_ is denoted by 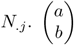 means the binomial coefficient.

NMI is another metric that represents the similarity of cluster sets *T* and *P*. It is formulated as:

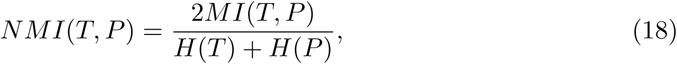

where *H*(·) denote the entropy of the cluster set and *MI*(·, ·) represents mutual information between cluster sets. Both ARI and NMI can be used to measure the consistency between the two data distributions. The domains of values of ARI and NMI are [−1, 1] and [0, 1], respectively. The larger the value, the more consistent the clustering results with the real situation.

#### Comparison of clustering performance

To evidence the effectiveness of CNLLRR, we perform clustering experiments on the scRNA-seq data. To make a fair comparison, CNLLRR and the other seven methods are provided with real cluster sets, even though the Corr itself can automatically estimate the number of clusters. In our experiment, ARI and NMI are adopted to evaluate the clustering performance. The experimental results are illustrated in Table 2.

**Table 2.**
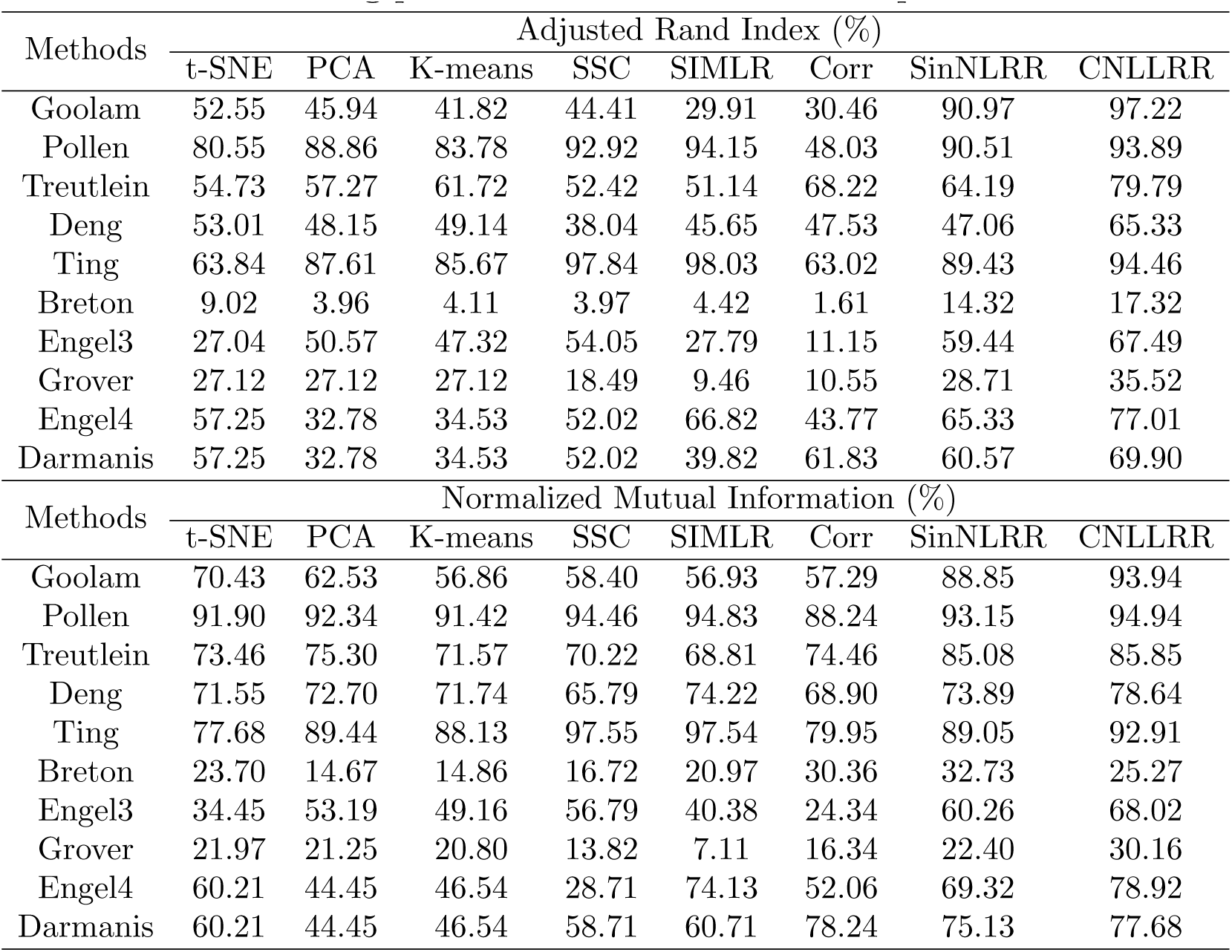
The clustering performance on the scRNA-seq data.

| Methods | Adjusted Rand Index (%) |  |  |  |  |  |  |  |
| --- | --- | --- | --- | --- | --- | --- | --- | --- |
|  | t-SNE | PCA | K-means | SSC | SIMLR | Corr | SinNLRR | CNLLRR |
| Goolam | 52.55 | 45.94 | 41.82 | 44.41 | 29.91 | 30.46 | 90.97 | 97.22 |
| Pollen | 80.55 | 88.86 | 83.78 | 92.92 | 94.15 | 48.03 | 90.51 | 93.89 |
| Treutlein | 54.73 | 57.27 | 61.72 | 52.42 | 51.14 | 68.22 | 64.19 | 79.79 |
| Deng | 53.01 | 48.15 | 49.14 | 38.04 | 45.65 | 47.53 | 47.06 | 65.33 |
| Ting | 63.84 | 87.61 | 85.67 | 97.84 | 98.03 | 63.02 | 89.43 | 94.46 |
| Breton | 9.02 | 3.96 | 4.11 | 3.97 | 4.42 | 1.61 | 14.32 | 17.32 |
| Engel3 | 27.04 | 50.57 | 47.32 | 54.05 | 27.79 | 11.15 | 59.44 | 67.49 |
| Grover | 27.12 | 27.12 | 27.12 | 18.49 | 9.46 | 10.55 | 28.71 | 35.52 |
| Engel4 | 57.25 | 32.78 | 34.53 | 52.02 | 66.82 | 43.77 | 65.33 | 77.01 |
| Darmanis | 57.25 | 32.78 | 34.53 | 52.02 | 39.82 | 61.83 | 60.57 | 69.90 |

| Methods | Normalized Mutual Information (%) |  |  |  |  |  |  |  |
| --- | --- | --- | --- | --- | --- | --- | --- | --- |
|  | t-SNE | PCA | K-means | SSC | SIMLR | Corr | SinNLRR | CNLLRR |
| Goolam | 70.43 | 62.53 | 56.86 | 58.40 | 56.93 | 57.29 | 88.85 | 93.94 |
| Pollen | 91.90 | 92.34 | 91.42 | 94.46 | 94.83 | 88.24 | 93.15 | 94.94 |
| Treutlein | 73.46 | 75.30 | 71.57 | 70.22 | 68.81 | 74.46 | 85.08 | 85.85 |
| Deng | 71.55 | 72.70 | 71.74 | 65.79 | 74.22 | 68.90 | 73.89 | 78.64 |
| Ting | 77.68 | 89.44 | 88.13 | 97.55 | 97.54 | 79.95 | 89.05 | 92.91 |
| Breton | 23.70 | 14.67 | 14.86 | 16.72 | 20.97 | 30.36 | 32.73 | 25.27 |
| Engel3 | 34.45 | 53.19 | 49.16 | 56.79 | 40.38 | 24.34 | 60.26 | 68.02 |
| Grover | 21.97 | 21.25 | 20.80 | 13.82 | 7.11 | 16.34 | 22.40 | 30.16 |
| Engel4 | 60.21 | 44.45 | 46.54 | 28.71 | 74.13 | 52.06 | 69.32 | 78.92 |
| Darmanis | 60.21 | 44.45 | 46.54 | 58.71 | 60.71 | 78.24 | 75.13 | 77.68 |

Based on Table 2, we can draw the following conclusions:

1. In the ten single-cell datasets, SinNLRR outperforms SIMLR and Corr by about 14% and 9% in terms of ARI and NMI. The CNLLRR is also superior to the SIMLR and Corr by around 23% and 13% on the metrics of ARI and NMI, respectively. The reason is that the LRR-based variants SinNLRR and CNLLRR capture the global structure of the data, while SIMLR and Corr only consider the similarity of the pair-wise data samples. This suggests that the LRR-based approach is more suitable for single-cell clustering due to its ability to capture complex relationships among samples.
2. From Table 2, we can observe that CNLLRR method exceeds SinNLRR method by approximately 9% and 4% on the AC and NMI metrics, respectively. There are mainly two reasons. First, using CLF instead of conventional norms on the error term in CNLLRR enable it with better robustness. Second, the graph regularization term applied in the proposed method explores local manifold information from data. On the account, CNLLRR has satisfactory clustering performance.
3. Basic simple clustering methods such as t-SNE, PCA, K-means, and SSC can achieve acceptable performances. For example, the average ARI score of SSC is 4% higher than of SIMLR and 12% higher than of Corr on ten single-cell data sets. This implies that improvements on traditional methods can not always promote clustering performance. For instance, the useful information may loss when naively modifying these methods, which can affect the clustering results.
4. It can be seen from Table 2 that our CNLLRR method can achieve best performance on most of datasets. Compared with other methods, the average scores of ARI and NMI of CNLLRR have approximately 9% and 4% improvements. We can conclude that our method is reasonable and effective in suppressing noise, outliers and preserving potential geometric information for the data.

In real-world applications, the number of clusters is usually unknown. To tackle this issue, we design an experiment to confirm the number of clusters in the scRNA-seq dataset. It is well known that SIMLR, Corr and SinNLRR are clustering methods proposed for the specific purpose of single-cell cluster analysis. Hence we use the three methods as comparison methods. In the experiment, we first construct the normalized Laplacian matrices **L**_*norm*_ which are based on the corresponding similarity matrices **S** obtained by each method. **L** is defined as

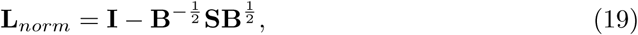

where **B** represents the diagonal matrix and the elements on its diagonal are **B**_*ii*_ = Σ_*j*_ **S**_*ij*_. **I** is the identity matrix. The eigengap [41] method is then applied on **L**_*norm*_, which can estimate the number of clusters by maximizing the eigenvalues gap |*λ*_*k*_ − *λ*_*k*−1_| of **L**_*norm*_. It should be mentioned that Corr decides the cluster number by itself automatically. Table 3 lists the comparison results of the four methods. We can see that CNLLRR concludes the same cluster number as the real cell types on the Treutlein, Ting and Engel3 datasets. In the remaining seven single-cell datasets, it obtains approximate cluster numbers as the number of real clusters. Although the performance of determining cluster numbers by all these methods is not satisfactory, the CNLLRR method is generally superior. It provides another perspective to validate the better clustering performance of our method.

**Table 3.** Comparison of the number of clusters estimated by the four methods.

| Datasets | Cell types | SIMLR | Corr | SinNLRR | CNLLRR |
| --- | --- | --- | --- | --- | --- |
| Goolam | 5 | 3 | 3 | 5 | 3 |
| Pollen | 11 | 19 | 3 | 10 | 10 |
| Treutlein | 5 | 12 | 3 | 4 | 5 |
| Deng | 7 | 6 | 4 | 5 | 5 |
| Ting | 5 | 51 | 5 | 5 | 5 |
| Breton | 4 | 2 | 3 | 6 | 2 |
| Engel3 | 3 | 2 | 3 | 5 | 3 |
| Grover | 2 | 4 | 6 | 4 | 4 |
| Engel4 | 4 | 2 | 2 | 6 | 2 |
| Darmanis | 8 | 2 | 6 | 10 | 7 |

### Visualization and gene markers

#### Visualize cells using t-SNE

In scRNA-seq data analysis, data visualization is also commonly used in identifying cell groups [42]. As a dimensionality reduction method, t-SNE has become a mainstream tool of visualization. According to [9], the modified t-SNE algorithm projects the input similarity matrix obtained by CNLLRR into a two-dimensional space, which shows the distinctions between cell subpopulations more intuitively and help biologists understand the diversity of cells more simply and directly.

Due to the limitation of space, we choose Engel4 and Darmanis datasets as the instances for data visualization to illustrate the performance of our method. Engel4 data are obtained from the thymus natural killer T (NTK) cells of female mice at 5 weeks [35]. It consists of 45 NKT0 cells, 46 NKT1 cells, 44 NKT17 cells and 68 NKT2 cells. Darmanis data contain 420 cells from adult and fetal cerebral cortical tissue [37]. Specifically, it has 110 fetal quiescent neurons, 25 fetal replicating neurons, 62 astrocytes, 131 neurons, 20 endothelial, 38 oligodendrocytes 16 microglia and 18 oligodendrocyte precursor (OPC) cells. Fig 8 shows the two-dimensional cell visualization results for t-SNE, SIMLR, SinNLRR, and CNLLRR on the Engel4 and Darmanis datasets. It can be seen from Fig 8(a) and 8(b) that t-SNE has the worst visualization effect. The clustering sets of the SIMLR method are more compact on the Engel4 and Darmanis datasets. This is because it requires the real cell labels to get the similarity matrix, whereas other methods do not. SIMLR always visualizes the same class of cells as different ones. For example, in Engel4 dataset, NTK0, NKT1, and NKT2 are not well distinguished. In constrast, as Fig 8 shows, the visualization result of CNLLRR is superior to SinNLRR and CNLLRR has better grouping results than most other methods. This above confirms the effectiveness of the CNLLRR method.

**Fig 8.**
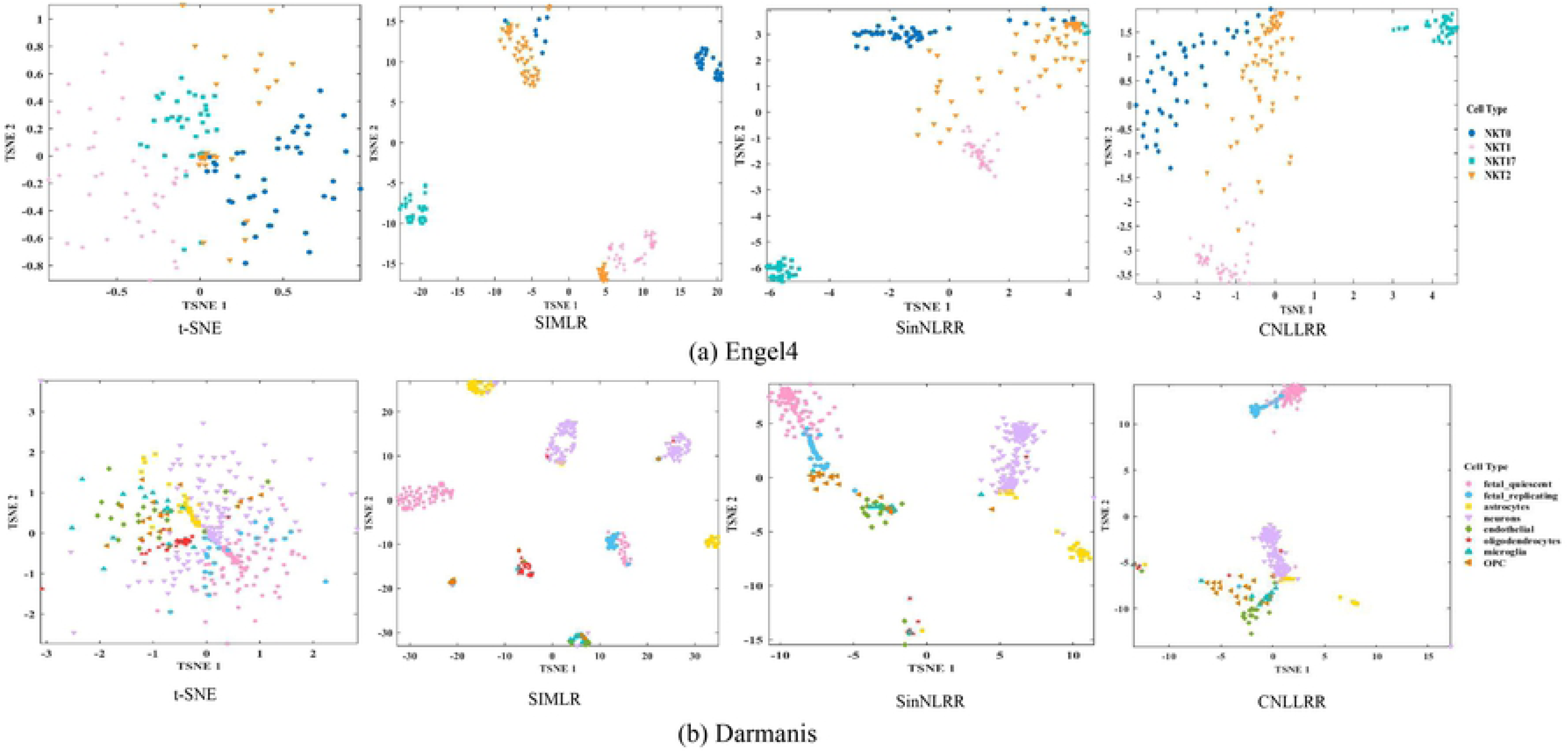
Comparison of cell visualization of the four methods. (a) Engel4 dataset. (b) Darmanis dataset.

#### Gene markers prioritization

The prioritization of gene markers has received intensive attentions since it was proposed. Cell gene markers which contain a wealth of valuable biological information are helpful in distinguishing cell subgroups and reveal cell complexity. The in-depth mining of biological information is of great significance in exploring cellular heterogeneity and life mechanisms. In our method, we use bootstrapped Laplacian score to identify gene markers on the similarity matrix obtained by CNLLRR [9]. The gene markers are then sorted in descending order corresponding to the importance of gene in distinguishing subgroups of cells.

Fig 9 demonstrates the top 10 gene markers in Engel4 and Darmanis datasets, and indicate their expression levels in different colors. The x-axis and y-axis represent the gene name and the cell type, respectively. As can be seen from Fig 9(a), the top 10 genes highly expressed in NKT2. REXO2, HMGB2 and CCT3 can be regarded as marker genes of NKT17, NKT1 and NKT0, respectively. REXO2 is a protein coding gene which may have a role in DNA repair, replication, and recombination, and in RNA processing and degradation. Therefore, the lack of REXO2 leads to a significant decrease in mitochondrial nucleic acid content [43]. HMGB2 encodes a member of non-histone chromosomal high mobility group protein family. The proteins of this family are chromatin-associated and distributed in the nucleus of higher eukaryotic cells ubiquitously [44]. The relevant pathways of CCT3 are cell cycle role of APC in cell cycle regulation and organelle biogenesis and maintenance. The protein encoded by this gene is a molecular chaperone that is a member of the chaperonin containing TCP1 complex (CCT), also known as the TCP1 ring complex (TRiC) [45]. In Fig 9(b), PLP1, TMEM144 and CLDND1 are the gene markers of oligodendrocytes. MALAT1 can be regarded as a gene marker of neurons. The gene markers of astrocytes are SLC1A2, SLC1A3, AQP4 and SPARCL1. MAP1B and TUBA1A are highly expressed on fetal quiescent neurons. The protein encoded by PLP1 may play a role in the compaction, stabilization, and maintenance of myelin sheaths, as well as in oligodendrocyte development and axonal survival. The protein encoded by AQP4 is the predominant aquaporin found in brain and has an important role in brain water homeostasis. Published articles further confirmed that PLP1 and AQP4 are the marker genes of oligodendrocytes and astrocytes, respectively [46]. To save space, we perform a brief analysis of some of the gene markers in the Engel4 and Darmanis datasets. The experiments and analysis provide new insights for researchers into the studies of NTK cells and cerebral cortical tissue.

**Fig 9.**
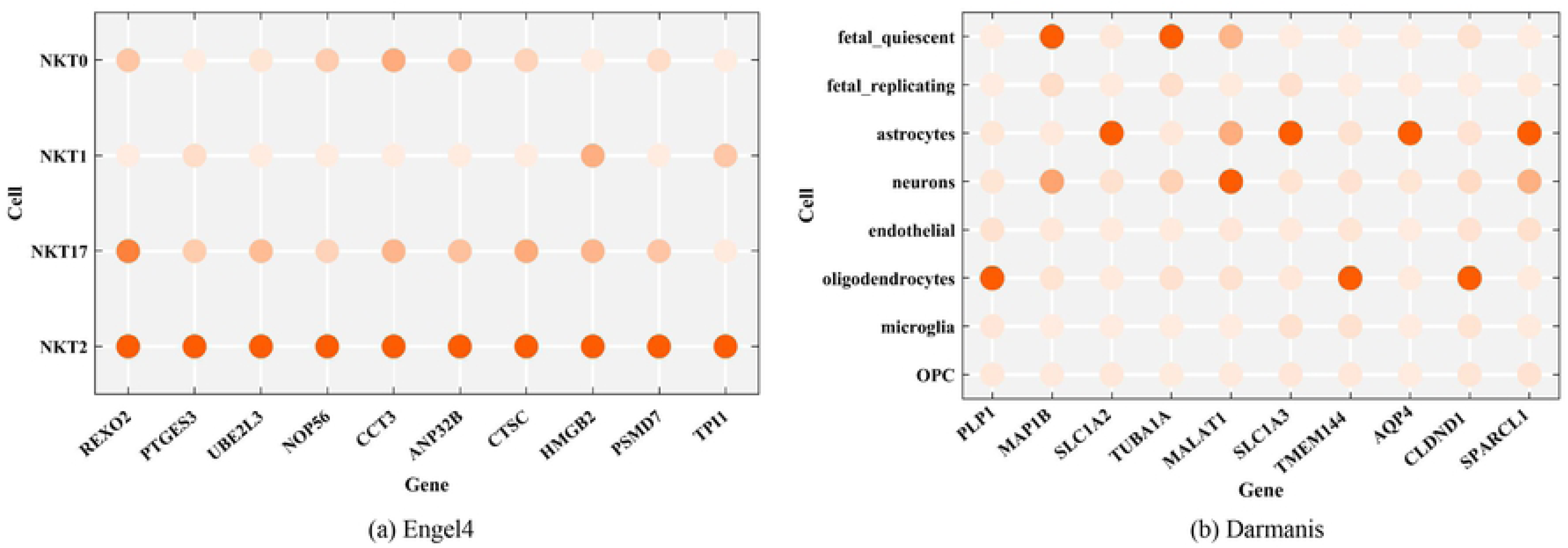
The top 10 gene markers of the two datasets. (a) Engel4 dataset. (b) Darmanis dataset.

## Conclusion

In this paper, we propose a robust LRR method called Cauchy non-negative Laplacian regularized LRR (CNLLRR) for scRNA-seq analysis. The proposed CNLLRR method uses CLF to reduce the negative impact of noise and outliers in clustering. Moreover, the graph regularization was applied to exploit the local manifold structure of data. Extensive experiments compared with other state-of-art methods on scRNA-seq data demonstrate the effectiveness of our proposed CNLLRR method. This method may provide a guiding for the next single-cell analysis and be applied to other relative fields.

For future work, we will make further efforts in model simplifying and application in larger dataset. In our model, the optimal solution is not easy to be obtained owing to the four free parameters. Another shortcoming of CNLLRR is the lack of flexibility when analyzing a large number of cells. Thus, we will continue to develop the Cauchy non-negative Laplacian regularized LRR method and promote its flexibility in the application of scRNA-seq analysis.

## Acknowledgments

This work was supported in part by the National Natural Science Foundation of China under Grant Nos. 61872220, 61873001, and 61972226.

